# Unique TMJ-specific transcriptomic signature and its medial layer: Implications in osteoarthritis

**DOI:** 10.1101/2025.05.18.654737

**Authors:** Rajnikant Dilip Raut, Chumki Choudhury, Amit Kumar Chakraborty, Harpreet Singh, Pushkar Mehra, Louis Gerstenfeld, Alejandro Almarza, Manish V. Bais

## Abstract

**Objectives:** Osteoarthritis (OA) is a debilitating joint disease that affects millions of people worldwide, with the temporomandibular joint (TMJ) and knee joint being prominently affected. Despite its prevalence, TMJ-OA remains understudied. This study aimed to investigate the transcriptional signature of the TMJ compared to that of the knee joint and to explore transcriptional differences in the medial and superficial layers of the TMJ-OA.

**Design:** Six-month-old C57BL/6J mice TMJ and knee samples were collected. Goat TMJ superficial and medial layer cartilage was separated and treated with IL-1β. All samples were subjected to bulk RNA sequencing followed by differential expression and gene set enrichment analysis.

**Results:** We identified 4,031 protein-coding genes differentially expressed in the TMJ compared to the knee, with significant enrichment of neuronal system genes and lower enrichment of innate immune system genes. Key osteoarthritis biomarkers such as *Mmp13, Postn*, and *Col1a1* were more highly expressed in the TMJ, indicating a higher vulnerability to OA development. IL-1β treatment in goat TMJ chondrocytes mimicked the natural TMJ-OA-like transcriptional changes and immune responses, which are also observed in the rabbit TMJ-OA model. This validated the *in vitro* goat TMJ-OA model. The IL-1β-treated goat TMJ medial cartilage layer was enriched in OA-associated transcription factors (TFs), senescence genes, and epigenetic regulators.

**Conclusion:** Our study demonstrated the unique transcriptomic signature of the TMJ compared with the knee joint, highlighting its vulnerability to OA and pain. These findings provide valuable insights into the molecular mechanisms of TMJ and offer a resource for potential therapeutic target selection for TMJ-OA treatment.

**Highlights:**

- Significant enrichment of neuronal system genes and lower enrichment of innate immune system genes in temporomandibular joint.
- Key osteoarthritis biomarkers such as *Mmp13, Postn*, and *Col1a1* have higher expression in temporomandibular joint, indicating a higher vulnerability to osteoarthritis development.
- Interleukin-1beta treatment in goat temporomandibular joint medial layer cartilage mimics natural temporomandibular joint osteoarthritis-like transcriptional changes and immune responses observed in rabbit temporomandibular joint osteoarthritis model.

## Introduction

Joint osteoarthritis (OA) leads to disability, pain, stiffness, cartilage degradation, and inflammation. The most affected joints include the temporomandibular joint (TMJ) and knee joints. Despite its prevalence, TMJ remains understudied, affecting 5-12% of the US population at an annual cost of US$ 4 billion (Scrivani et al., 2008; Velly et al., 2013). The TMJ functions through rotation and translation during chewing, breathing, and speaking(Y.-H. Lee et al., 2020). The prevalence of TMJ-OA is higher in females compared than in knee OA and presents a severe pathology. The TMJ, which is responsible for jaw movements, and the knee joint, which is crucial for locomotion, exhibit unique biomechanical environments influencing OA progression(Bielajew et al., 2021). Clinically, TMJ-OA is distinct from knee OA(Vos et al., 2014). Understanding the specific molecular pathways and transcriptional changes in the TMJ compared to knee OA is essential for developing targeted therapies. The TMJ has secondary cartilage compared to the knee joint, suggesting differences in the superficial and medial layers of the condylar cartilage.

Molecular cues promote mesenchymal stem cells or chondroprogenitors to regenerate the TMJ and knee joint articular cartilage(Embree et al., 2016; Jiang & Tuan, 2015; M.-S. Lee et al., 2021; Wei et al., 2021; Worthley et al., 2015). The TMJ contains both fibrocartilage and mandibular condylar cartilage, which distinguishes it from the articular cartilage of the limbs and the cartilage in the cranial base(Shibata et al., 1997). TMJ cartilage expresses specific proteins and has structural differences compared with other joints(Hinton, 2014). Structurally, there are different layers of chondrocytes in TMJ cartilage(Wadhwa & Kapila, 2008).

Considering TMJ’s molecular uniqueness and translational applications of the TMJ, we explored intra-TMJ cartilage variations using goat TMJ cartilage. The TMJ cartilage is composed of two layers: the superficial layer cartilage (SLC) and middle layer cartilage (MLC). An interesting study by *Tosa et al*., delineated the transcriptomic profiles of SZ and CC in New Zealand white rabbit(Tosa et al., 2023). However, this study largely omitted the variability in inflammatory processes and OA-specific gene expression. We segregated SLC and MLC from goat TMJ cartilage and performed bulk RNA sequencing to determine their transcriptomic profiles and their utility in TMJ-OA.

Our earlier studies on the evaluation of LOXL2 as a therapeutic target in TMJ-OA therapy suggested that there could be distinct mechanisms and pathways within the TMJ cartilage compared with the knee(Alshenibr et al., 2017; M. Tashkandi et al., 2019; M. M. Tashkandi et al., 2020). This study characterizes TMJ transcriptional differences from the knee joint, differences in the superficial and medial layers of the TMJ, and responses to OA-related factors using RNA sequencing. Specifically, we evaluated mouse TMJ transcriptome differences compared to those in the knee joint and characterized them using *in vitro* goat chondrocytes. A comparison of goat TMJ chondrocytes *in vitro* model with rabbit *in vivo* models revealed unique signaling pathways. By elucidating the mechanisms of OA in these joints, we aimed to provide insights into effective and personalized TMJ-OA-specific targets. The molecular pathways and gene expression profiles of TMJ cartilage could serve as a reference for developing novel therapies for TMJ-OA treatment(Ma et al., 2025).

## Materials and methods

### Animal studies: TMJ and knee isolation

The animal experiments have been approved by Institutional Animal Care and Use Committee (IACUC AN-15387) of Boston University. Six-month-old C57BL/6J mice were obtained from Jackson laboratory (#664). Mice were euthanized by CO_2_ following approved euthanasia guidelines. Knee joints were collected by separating the entire leg by cutting from the proximal epiphysis of the femur; cleared of hairs, muscles and fat tissue by scissor and scalples, and a final rubbing with gauze. Complete knee integrity was maintained and joints were collected by cutting at femoral distal metaphysis and proximal tibia. We collected the TMJ condyles from ascending ramus using surgical scissor. Thereafter, samples were snap frozen, crushed, subjected to RNA extraction with TRIzol, and sent for RNA sequencing at NOVOGENE Sacramento, California (n=4 TMJs and n=4 Knee joints).

### Goat TMJ chondrocyte cell culture

Fresh goat TMJ samples were obtained from local abattoirs (age 9 months, female) in the Almarza laboratory and grown as a bilogical replicates and various preparations. The superficial layer cartilage (SLC) of the condylar fibrocartilage was separated from the medial layer cartilage (MLC) by digesting it for an hour using 200U/mL collagenase type II, in accordance with the technique developed by the Almarza lab. The SLC was digested for an additional hour before being peeled, chopped, and digested for three more hours in 200 U/mL collagenase type II (Worthington). Using a scalpel blade, MCL was extracted from TMJ bone and digested in 200 U/mL collagenase type II (Worthington) at 37°C and 5% CO2 under mechanical stirring (rocker, approximately 0.4 Hz). Subsequently, both cell types were plated in culture media in the respective groups for a further twenty-four hours. The cells were then sent to Boston University after freezing in dimethyl sulfoxide (DMSO). Standard culture conditions (37°C, 5% CO2) were used for both SLC and MLC, using DMEM supplemented with 10% FBS. Both SLC and MLC were grown in Dulbecco’s Modified Eagle Medium (DMEM) containing 10% FBS under normal conditions (37°C, 5% CO2). Cells were used at passages 3–5. The treatment consisted of a vehicle and IL-1β (10ng). The IL-1β therapy lasted for an hour. Following treatment, cells were harvested for RNA sequencing and downstream analysis.

### RNA isolation and sequencing

Total RNA was isolated from mice and goat samples (*Please see animal studies methods*) using TRIzol reagent (Qiagen) according to the manufacturer’s instructions. Extracted RNA samples were sequenced on Novogene (Sacramento, California, United States). All samples were qualitatively tested prior to library construction. Only samples that passed the quality check based on the RNA Integrity Number (RIN) were used. The Illumina high-throughput sequencing platform was used for paired-end RNA sequencing.

### Bulk RNA-seq data processing and bioinformatics analyses

Raw FASTQ sequencing reads were pre-processed or trimmed using an in-house Novogene Perl script. The filtered data were aligned to the reference genomes of Capra hircus (ncbi_capra_hircus_gcf_001704415_2_ars1_2) and Mus musculus (mm10). Raw reads corresponding to each gene were counted using Feature Counts (v1.5.0-p3). The DESeq2 software in R/Bioconductor was used to determine the differential gene expression (DGE). Gene set enrichment analysis (GSEA-v4.3.2) software was used to conduct functional enrichment analysis to assess implicated biological processes or pathways.

Publicly available rabbit TMJ bulk RNA seq data was obtained from GSE232867. Here, the authors have performed a comparative gene expression profiling analysis for control and post-traumatic TMJ-OA model in superficial zone cells and condylar cartilage cells.

### Scanning Electron Microscopy (SEM)

To prepare for observation under scanning electron microscope, the cultured goat cells on glass were fixed with 2.5% glutaraldehyde fixation overnight, and 1% osmium tetroxide for 30 mins post fixation, treated with serial dehydration using ethanol from 25%, 50%, 75% to 100%, then critical point dried with carbon dioxide critical point dryer (Samdri PVT-3B). The dried samples were mounted on an aluminum SEM stub and sputter-coated with gold/palladium (75/25) using a sputter coater (Hummer V, Anatech) to improve conductivity. The cell morphology and organization were observed using field-emission SEM (SU6600; Hitachi) at an accelerating voltage of 15 kV at different magnifications.

### Statistical analysis

The Complex Heatmap package in R was used to create the heat maps. Using R’s ggplot2 package in R, volcano plots were created for genes with differential expression. GraphPad PRISM (v10.1.0) was used to create box/bar plots. For DGE analysis, DESeq2 uses wald test to caluculate the p-values and adjusted for multiple testing using the Benjamini-Hochberg method to get the adjusted p-values.

## Results

### TMJ has a unique transcriptomics signature as compared to knee joint

Molecular variations in the TMJ compared to the knee are not known. Thus, we first evaluated transcriptomic differences using RNA sequencing, followed by differential gene expression (DGE) and gene set enrichment analysis (GSEA). A total of 4031 protein-coding genes were differentially expressed in the TMJ, which were selected based on an adjusted p-value < 0.05, using the Benjamini-Hochberg method (n=4 mice/group) (Fig. 1A). Almost equal numbers of genes were highly expressed in both the TMJ (n=2045) and the knee (n=1986). GSEA revealed the enrichment status (positive or negative) of gene sets related to the neuronal system, innate immune system, collagen degradation, and extracellular matrix (ECM) degradation in the TMJ (Fig. 1B, C). It appears that most genes in the innate immune system are expressed at low levels in the TMJ compared to the knee because of the higher expression of neuronal system genes to balance the inflammatory pain response in the TMJ.

**Fig. 1:**
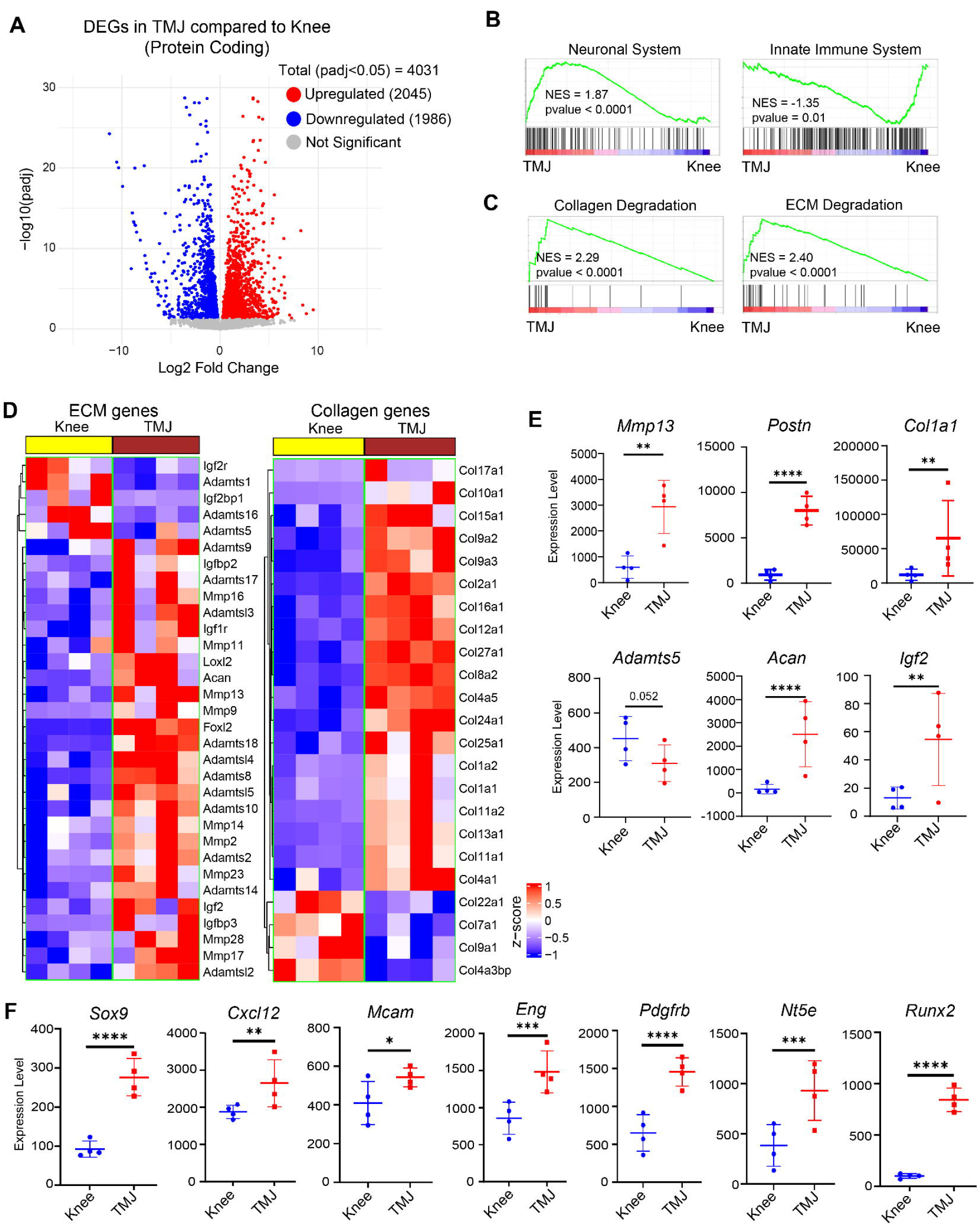
Transcriptomic variations in the TMJ compared to knee joint. **A)** Differentially expressed protein coding genes between TMJ and Knee; Selected based on adjusted p-value < 0.05 (Benjamini-Hochberg method) (n=4 mice/group). **B-C)** Gene set enrichment analysis (GSEA) showing enrichment status (positive or negative) of gene sets related to neuronal system, innate immune system, collagen degradation, and ECM degradation in the TMJ; NES: Normalized Enrichment Score. **D)** Heatmap representation of top differentially expressed extracellular matrix (ECM) genes between knee and TMJ; Red: higher expression, Blue: lower expression. **E)** Box plots representing expression of OA associated genes between knee and TMJ. **F)** Box plots representing expression of non-osteogenic and osteogenic regeneration genes between knee and TMJ. *p<0.05, **p<0.01, ***p<0.001, ****p<0.0001. P values were calculated using the default Wald test in DESeq2. Error bars represent Mean with standard deviation (SD).

A heatmap representation highlighted the top differentially expressed extracellular matrix (ECM) genes between the knee and TMJ, with red indicating higher expression and blue indicating lower expression (Fig. 1D). Similarly, most of the key collagen factors were more highly expressed in the TMJ than in the knee (Fig. 1D). Studies have suggested that *Mmp13, Postn, Col1a1*, and *Adamts5* are key biomarkers of osteoarthritis when overexpressed. Surprisingly, the TMJ showed higher expression of these OA biomarkers, but not *Adamts5*, rendering this constantly locomotive joint vulnerable to OA development (Fig. 1E). Moreover, *Acan* and *Igf2* were highly expressed in the TMJ (Fig. 1E). Next, we analyzed the expression of important cartilage regeneration factors during cartilage damage and OA. Specifically, we observed higher expression of non-osteogenic genes such as *Sox9, Cxcl12* (SDF1), *Mcam* (CD146), *Eng* (CD105), *Pdgfrb*, and osteogenic genes such as *Nt5e* (CD73) and *Runx2* in TMJ (Fig. 1F). These data reveal a significant molecular difference between the TMJ and the knee joint and suggest that the TMJ is highly vulnerable to OA and pain.

### TMJ medial layer has a unique transcriptional signature as compared to the superficial layer

To determine the molecular uniqueness of the TMJ, we explored the intra-TMJ cartilage molecular variations using global transcriptomic profiling. We extracted both the superficial layer cartilage (SLC) and medial layer cartilage (MLC) from goat TMJ condyle cartilage using the protocol described in the Methods section and subjected them to bulk RNA sequencing (n=4/group). A total of 695 genes in SLC and 901 genes in MLC were highly expressed based on the stringent selection criteria of an adjusted p-value below 0.05 and log2FC above 1 (Fig. 2A). GSEA suggested that gene sets related to ECM proteoglycans and ECM degradation were highly enriched in MLC compared to those in SLC, making it susceptible to TMJ-OA development (Fig. 2B). This was complemented by higher enrichment of gene sets related to collagen formation and degradation in MLC (Fig. 2B).

**Fig. 2:**
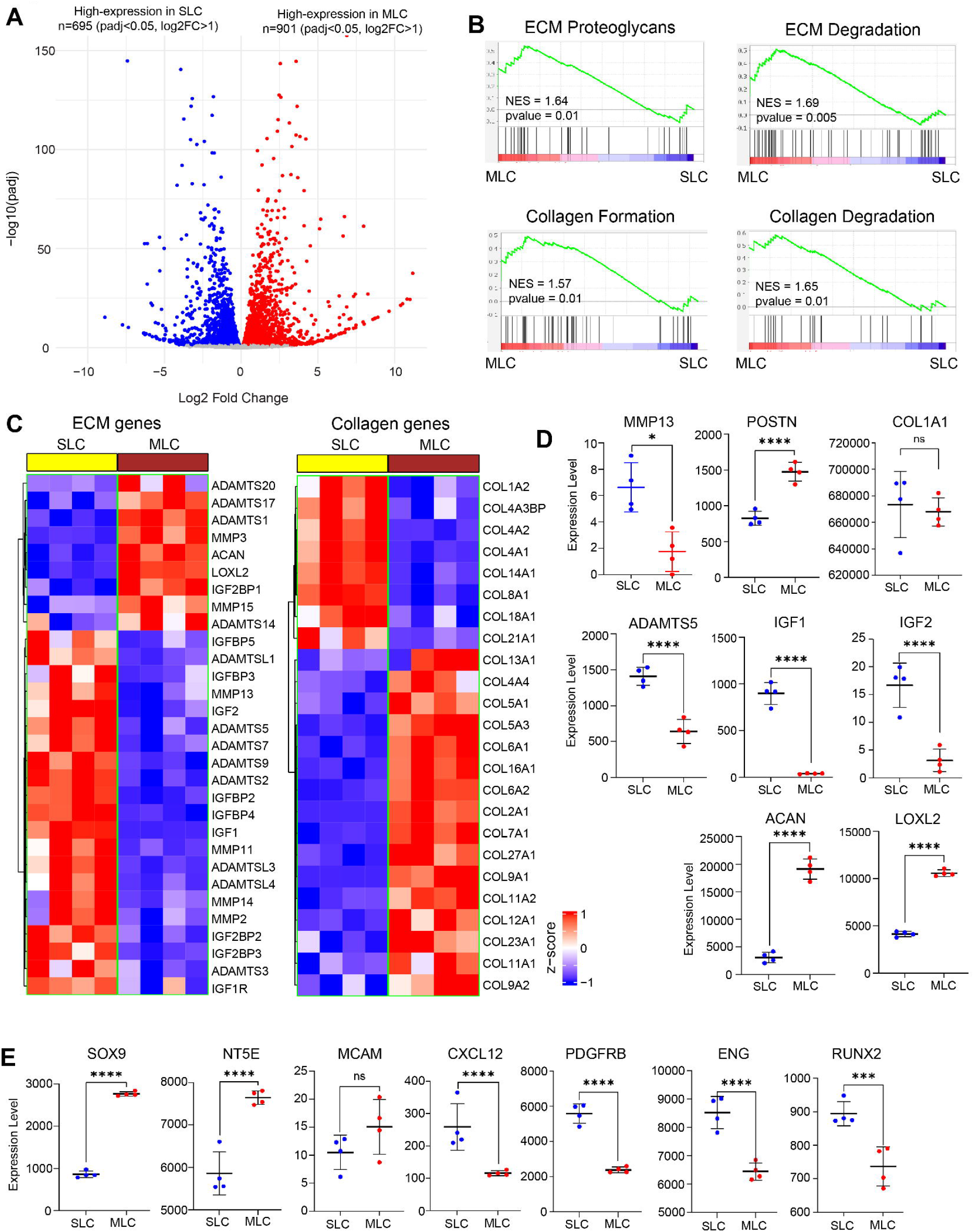
TMJ superficial layer cells (SLC) are transcriptionally distinct than medial cartilage layer cells (MLC). **A)** Differentially expressed genes between SLC and MLC; Selected based on adjusted p-value < 0.05 (Benjamini-Hochberg method) and log2 fold change > 1 (n=4/group). **B)** GSEA showing enrichment status of gene sets related to ECM proteoglycans, ECM degradation, collagen formation, and collagen degradation in the goat TMJ SLC and MLC; NES: Normalized Enrichment Score. **C)** Heatmap representation of top differentially expressed extracellular matrix (ECM) genes between SLC and MLC; Red: higher expression, Blue: lower expression. **D)** Box plots representing expression of OA associated genes, growth factors, and ECM genes between SLC and MLC. **E)** Box plots representing expression of non-osteogenic and osteogenic regeneration genes between knee and TMJ. *p<0.05, **p<0.01, ***p<0.001, ****p<0.0001. P values were calculated using the default Wald test in DESeq2. Error bars represent Mean with SD.

Although the ECM and collagen degradation gene sets were highly enriched in MLC, heatmap analysis showed that most ECM and collagen degradation genes had lower expression levels in MLC than in SLC, excluding the OA-specific gene MMP3, a collagen type 2 degrading enzyme (Fig. 2C). Interestingly, most of the collagen genes were highly expressed in MLC, including COL2A1, an essential factor for the structure and function of cartilage (Fig. 2C). Additionally, the expression of key OA factors MMP13 and ADAMTS5 was lower, whereas POSTN was highly expressed in MLC, and the expression of COL1A1 was similar in both MLC and SLC (Fig. 2D). Growth factors such as IGF1 and IGF2 were highly expressed in SLC but not in MLC, whereas ACAN and collagen crosslinking factor LOXL2 were highly expressed in MLC (Fig. 2D). Expression analysis of regeneration factors revealed that MLC was the hub for SOX9 and NT5E, whereas CXCL12, PDGFRB, ENG, and RUNX2 were highly expressed in SLC (Fig. 2E). There was no significant difference in the MCAM expression between the SLC and MLC groups. As both layers expressed non-osteogenic and osteogenic markers, it can be concluded that SLC and MLC played a mutual role during the cartilage regeneration process by maintaining the dynamics of osteogenic and non-osteogenic cellular composition in the TMJ cartilage.

### TMJ medial layer is highly vulnerable to IL-1β treatment and mimics natural TMJ-OA-like changes

Overexpression of IL-1β is a major biomarker of osteoarthritis development. We examined the effects of IL-1β treatment on superficial layer cartilage (SLC) and medial layer cartilage (MLC). To visualize the phenotypic effects of IL-1β treatment on both SLC and MLC, chondrocytes were incubated with IL-1β for 1 h and subjected to scanning electron microscopy (SEM). As observed from the SEM images, IL-1β treatment resulted in degradation of the extracellular matrix-like (ECM-L) structure around the chondrocytes (Fig. 3A, B). Specifically, the ECM-L structure around the SLC was degraded and the overall cellular phenotype was similar to that of apoptotic cells previously observed by SEM (Fig. 3A). Importantly, this effect was more severe in IL-1β treated MLC because most of the ECM-L structure was completely degraded (Fig. 3B). This analysis suggests that MLCs are highly susceptible to IL-1β treatment and IL-1β treated MLC could be considered a sustainable model system for in vitro molecular studies in OA, as ECM degradation in cartilage is a hallmark of OA.

**Fig. 3:**
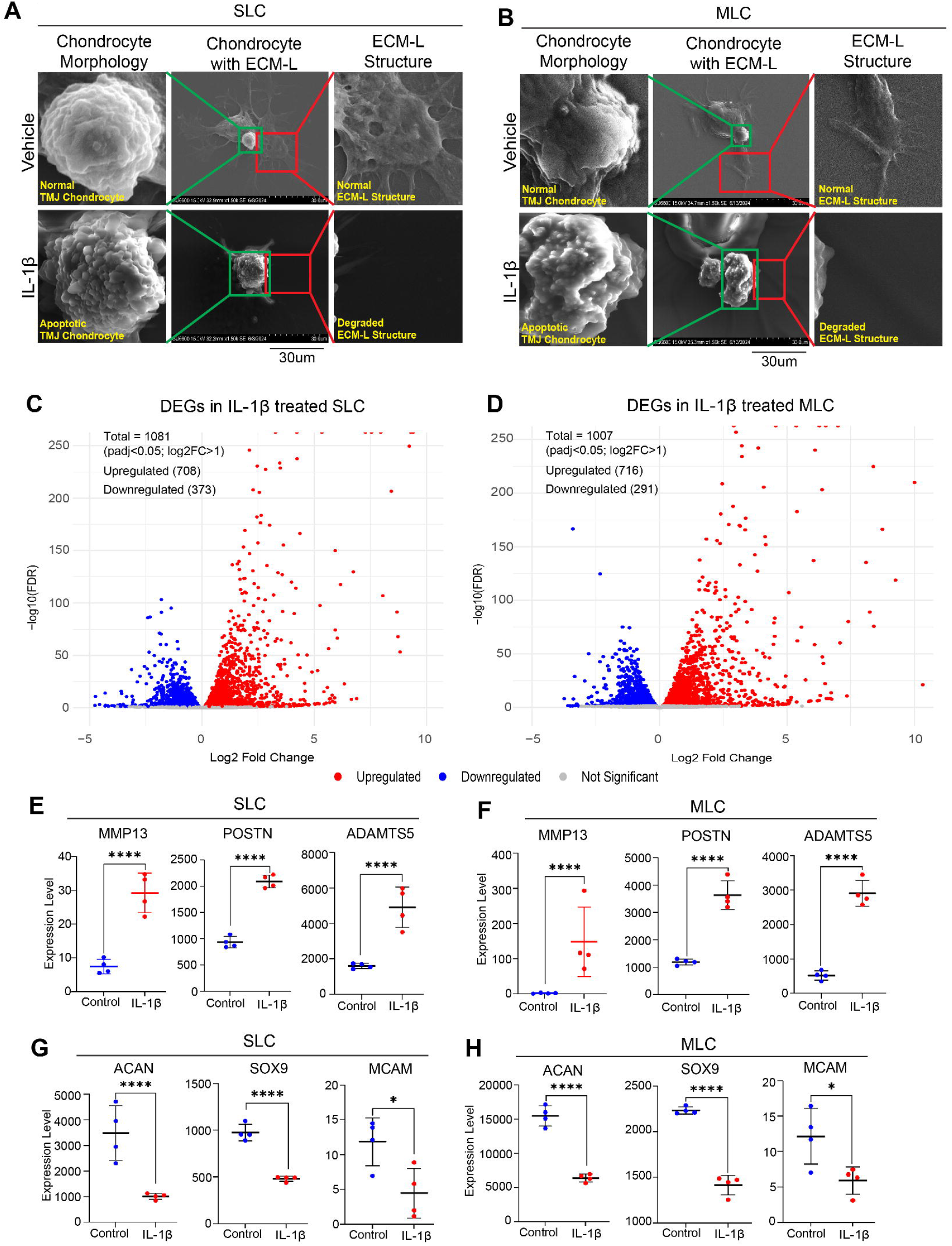
IL-1β treatment in the goat TMJ cartilage (SLC and MLC) mimics natural TMJ-OA-like molecular changes and immune response. **A)** Scanning electron microscopy (SEM) images of SLC along with its ECM-L structure at 1500x resolution; ECM-L structure is degraded around SLC and turned apoptotic upon IL-1β treatment. **B)** SEM images of MLC along with its ECM-L structure at 1500x resolution; ECM-L structure is completely degraded around MLC and turns apoptotic upon IL-1β treatment. **C)** Differentially expressed genes in SLC IL-1β treatment; Selected based on adjusted p-value < 0.05 (Benjamini-Hochberg method) and log2 fold change > 1 (n=4/group). **D)** Differentially expressed genes in MLC IL-1β treatment; Selected based on adjusted p-value < 0.05 (Benjamini-Hochberg method) and log2 fold change > 1 (n=4/group). **E-F)** Bar plots showing increased mRNA expression of OA genes MMP13, POSTN, and ADAMTS5 after IL-1β treatment in SLC and MLC. **G-H)** Bar plots showing decreased expression of ECM (ACAN) and non-osteogenic regeneration genes (SOX9 and MCAM) after IL-1β treatment in SLC and MLC. *p<0.05, **p<0.01, ***p<0.001, ****p<0.0001. P values were calculated using the default Wald test in DESeq2. Error bars represent Mean with SD.

To determine the transcriptomic profiles of SLC and MLC after IL-1β treatment, we performed bulk RNA sequencing followed by differential gene expression (DGE) analysis (n=4/condition). We observed significant overexpression of genes in IL-1β treated samples based on stringent selection criteria of log2 fold change above 1 and adjusted p-value below 0.05. In IL-1β treated SLC, a total of 1081 genes were differentially expressed, with around 708 genes upregulated and 373 genes downregulated (Fig. 3C). In IL-1β treated MLC, a total of 1007 genes were differentially expressed, with approximately 716 genes upregulated and 391 genes downregulated (Fig. 3D). Next, we analyzed the expression of key OA development genes such as MMP13, POSTN, and ADAMTS5, along with the cartilage regeneration factors SOX9, MCAM, and ACAN. As expected, all three OA genes were highly overexpressed in IL-1β–treated SLCs and MLCs (Fig. 3E, F). Interestingly, the expression levels of these genes were much higher in IL-1β–treated MLCs than in SLCs. This transcriptomic profile explains why IL-1β mediated degradation under SEM was significantly more severe in the MLCs than in the SLCs. Additionally, cartilage regeneration genes such as SOX9, MCAM, and ACAN, which are key components of the cartilage extracellular matrix for normal functional homeostasis, were also downregulated in IL-1β treated samples (Fig. 3G, H). Taken together, the SEM results in agreement with the transcriptomic profile of IL-1β treated SLC and MLC suggest that MLCs are highly prone to degradation and are major contributors to OA development.

### IL-1β induced OA-like changes in MLC correlate with the Rabbit TMJ-OA model

To validate our in vitro goat MLC model, we compared IL-1β induced changes in the in vitro goat TMJ-OA model with rabbit TMJ-OA models. We re-analyzed the bulk RNA-seq data of rabbit TMJ-OA SZ and CC from Tosa et al.,(Tosa et al., 2023), performing differential gene expression (DGE) analysis followed by gene set enrichment analysis (GSEA) to identify dysregulated genes and signaling pathways. In our study, SZ was represented as SLC and CC as MLC to maintain coherence, as both layers originated from the same TMJ cartilage region in rabbits and goats. IL-1β treated goat SLC showed consistent up- and down-regulation of genes; however, we found a large number of overexpressed genes in MLC, which also correlated with the IL-1β treated goat MLC profile (Fig. 4A, B). Furthermore, GSEA analysis of rabbit TMJ-OA MLC showed that chondrocyte apoptosis via the cell killing pathway was highly enriched (Fig. 4C). Inflammatory immune response-related gene sets were strongly enriched, including mast cell and macrophage activation, positive regulation of cytokine production and signaling, neuroinflammatory responses, and gene sets involved in the elevated production of IL-1 and IL-6 (Fig. 4C). Interestingly, all these gene sets were also highly enriched in IL-1β treated goat MLC, recapitulating the inflammatory transcriptomic profile of in vivo rabbit TMJ-OA MLC in the in vitro goat TMJ-OA model (Fig. 4D).

**Fig. 4:**
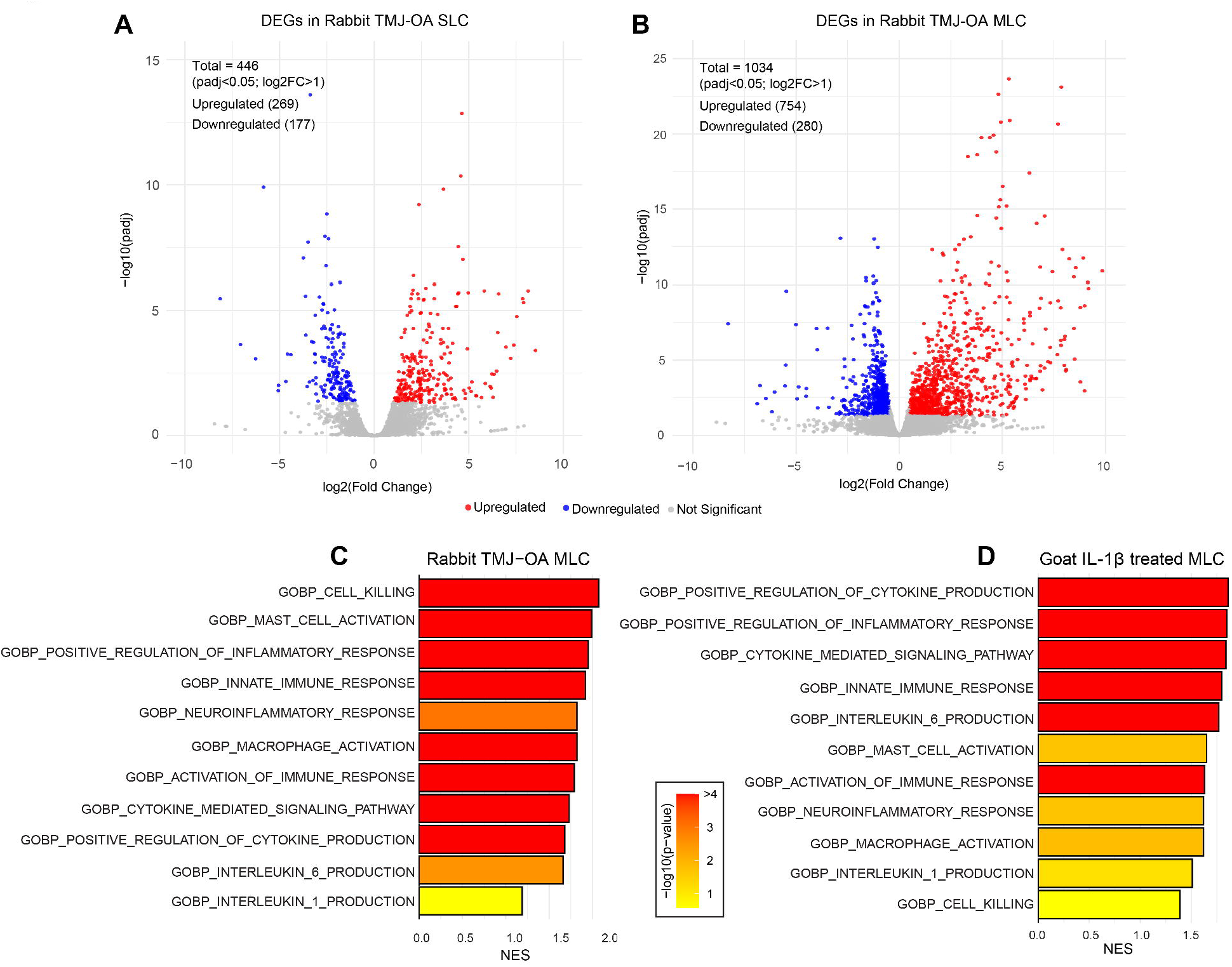
IL-1β Induced Goat MLC has similarity with rabbit TMJ-OA model. **A-B)** Re-analysis of differentially expressed genes in superficial zone (here, SLC) and condylar cartilage (here, MLC) of surgically induced rabbit TMJ-OA model; Selected based on adjusted p-value < 0.05 (Benjamini-Hochberg method) and log2 fold change > 1 (n=3/group). **C)** GSEA showing enrichment status of gene sets related to cell killing and inflammatory pathways enrichment in the MLC of rabbit TMJ-OA model. D) GSEA showing enrichment status of gene sets related to cell killing and inflammatory pathways enrichment in IL-1β treated MLCs, which correlates with rabbit TMJ-OA MLC like inflammatory processes. NES: Normalized Enrichment Score.

**Fig. 5.**
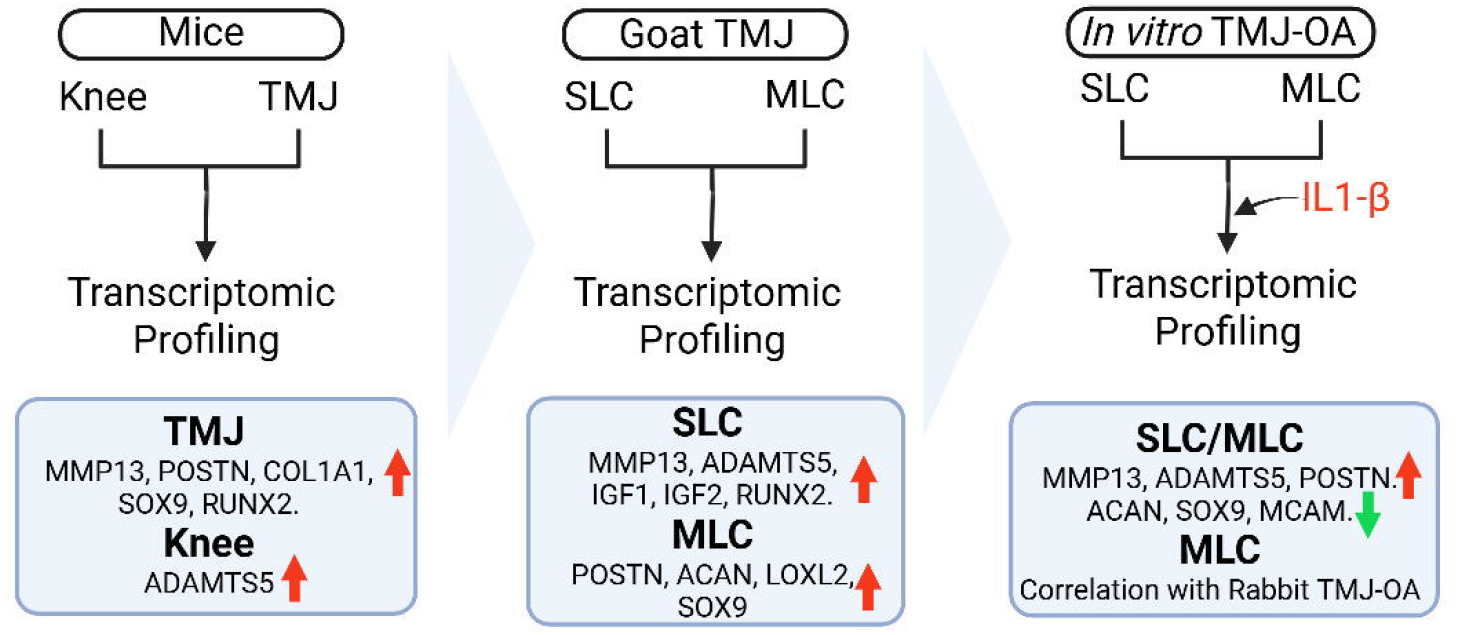
Graphical abstract illustrating findings of this study

### IL-1β treatment activates OA associated transcription factors, senescence genes, and epigenetic regulators in goat TMJ condylar cartilage layer

To identify TMJ-OA-specific genes expressed in our goat TMJ-OA MLC model, we explored published clinical knee-OA transcription factors (TFs)(Swahn et al., 2023) as clinical TMJ-OA-specific genes were not identified. Our study found that IL-1β treated goat MLC, which showed OA-like molecular changes, had significant overexpression of most of these OA-associated TFs (Table 1).

**Table 1.** Gene expression of OA associated TFs, senescence genes, and epigenetic regulators in IL-1β treated MLCs.

|  | Gene | log2FoldChange | pvalue | padj |
| --- | --- | --- | --- | --- |
| **Transcription Factors** | OLFML2B | 4.128345016 | 6.81E-50 | 6.33E-48 |
|  | SERPINE2 | 3.227229635 | 4.34E-248 | 7.80E-245 |
|  | SFRP4 | 3.154713341 | 2.08E-136 | 9.10E-134 |
|  | NPR3 | 2.989267548 | 1.54E-47 | 1.31E-45 |
|  | IGFBP3 | 2.503814145 | 6.88E-37 | 4.17E-35 |
|  | ADAMTS7 | 2.297229768 | 1.38E-64 | 1.99E-62 |
|  | NOV | 2.213349365 | 1.41E-41 | 9.89E-40 |
|  | TREM1 | 2.025665131 | 5.60E-53 | 5.78E-51 |
|  | WNT5A | 1.838856848 | 1.59E-160 | 8.89E-158 |
|  | TNFAIP6 | 1.794384692 | 1.71E-18 | 4.51E-17 |
|  | ITGB8 | 1.789700247 | 8.88E-52 | 8.76E-50 |
|  | CACNA1A | 1.502783932 | 2.26E-09 | 3.06E-08 |
|  | TNFSF15 | 1.327715561 | 8.50E-06 | 7.32E-05 |
|  | SEMA3C | 1.316547207 | 6.46E-33 | 3.39E-31 |
|  | TLL1 | 1.268895132 | 1.48E-21 | 4.60E-20 |
|  | EPDR1 | 1.193671159 | 5.00E-46 | 3.97E-44 |
|  | SULF1 | 1.112470577 | 1.42E-62 | 1.96E-60 |
|  | ADAMTS2 | 1.034519987 | 1.24E-55 | 1.38E-53 |
| **Senescence genes** | CSF2 | 6.549657173 | 4.03E-10 | 5.85E-09 |
|  | ICAM1 | 6.100384789 | 5.23E-244 | 7.70E-241 |
|  | CCL5 | 5.729394373 | 4.98E-28 | 2.14E-26 |
|  | IL1A | 4.915758936 | 6.91E-22 | 2.17E-20 |
|  | MMP12 | 4.413820924 | 1.19E-09 | 1.64E-08 |
|  | EREG | 2.761738562 | 6.42E-79 | 1.32E-76 |
|  | PGF | 1.902925586 | 0.01042564 | 0.041503448 |
|  | IGFBP4 | 1.646813552 | 1.25E-73 | 2.17E-71 |
|  | CXCL16 | 1.406750044 | 6.16E-35 | 3.52E-33 |
|  | PLAT | 1.263407961 | 3.03E-27 | 1.25E-25 |
| **Epigenetic Regulators** | ARRB1 | 2.387682523 | 2.26E-11 | 3.67E-10 |
|  | ARID3A | 1.880751479 | 2.47E-29 | 1.10E-27 |
|  | PRDM1 | 1.862526076 | 2.26E-05 | 0.000179882 |
|  | ARID5A | 1.142968943 | 2.03E-12 | 3.62E-11 |
|  | GTF3C1 | 1.133709667 | 6.65E-56 | 7.63E-54 |

These include TF such as ADAMTS7, which mediates the degradation of cartilage oligomeric matrix protein (COMP), a key component of cartilage, and forms a positive feedback loop with TNFα, contributing to inflammation and OA pathogenesis(Liu et al., 2006; Zhang et al., 2015). The NOV (CCN3) gene is associated with chondrocyte senescence and cartilage degradation via induction of p53 and p21(Kuwahara et al., 2020).

Cellular senescence is a major hallmark of OA progression. Therefore, we analyzed the expression of senescence- and senescence-associated secretory factors(Swahn et al., 2023) (Table 1). These factors were markedly overexpressed after IL-1β treatment in goat MLC, confirming the specificity of this *in vitro* model for TMJ-OA studies. Genes such as CSF2 are involved in the development of pain-like behavior in OA models(Shin et al., 2023). ICAM1 expression has been found to be upregulated in the synovium of OA patients(Law et al., 2020). Moreover, CXCL16 also exerts inflammatory effects on synovium(van der Voort et al., 2005).

In addition, we observed the overexpression of previously unidentified epigenetic regulators that contribute to OA development and progression (Table 1). ARID3A has not been studied in OA; however, it is strongly associated with IFNγ production(Ratliff et al., 2019). ARID5A stabilizes IL6 mRNA, thus contributing to inflammation and OA pathogenesis(Masuda et al., 2013). Taken together, IL1B treatment in goat MLC favors hyperactivation of TFs, senescence genes, and epigenetic regulators that are crucial in OA pathophysiology.

## Discussion

Although phenotypic differences and histological changes in the TMJ have been characterized, molecular changes are not completely known. This study aimed to investigate the transcriptional signature of mouse TMJ compared to the knee joint, explore differences in TMJ medial and superficial layers, and evaluate *in vitro* goat TMJ-OA model, which could be exploited for translational studies as well as to predict the TMJ-OA-specific gene signature.

Transcriptomic analysis of the TMJ compared with the knee joint revealed significant molecular differences that highlight the unique characteristics of the TMJ. Our RNA sequencing and DGE analysis identified 4031 protein-coding genes that were differentially expressed in the TMJ, with almost equal numbers of genes highly expressed in both the TMJ and the knee. Furthermore, GSEA revealed enrichment of gene sets related to the neuronal system, innate immune system, collagen degradation, and ECM degradation in the TMJ. Notably, the TMJ exhibited a lower expression of innate immune system genes, potentially because of the higher expression of neuronal system genes that may balance the inflammatory pain response.

Heatmap analysis highlighted the top ECM genes differentially expressed between the knee and TMJ, with key collagen factors showing higher expression in the TMJ. Interestingly, biomarkers of osteoarthritis (OA), such as *Mmp13, Postn*, and *Col1a1*, excluding *Adamts5*, were more highly expressed in the TMJ, suggesting a higher vulnerability to OA development. Additionally, cartilage regeneration factors such as *Acan* and *Igf2* were highly expressed in the TMJ, indicating a complex interplay between degeneration and regeneration processes.

Intra-TMJ molecular variations were further explored through the global transcriptomic profiling of SLC and MLC. Our analysis revealed that MLC is more susceptible to TMJ-OA development, with higher enrichment of gene sets related to ECM proteoglycans and collagen formation and degradation. Despite enrichment, most ECM and collagen degradation genes had lower expression in MLC than in SLC, except for the OA-specific gene MMP3. The expression of key OA factors varied between layers, with POSTN being highly expressed in MLC and similar levels of COL1A1 in both layers. Growth factors such as IGF1 and IGF2 were differentially expressed, with IGF1 being highly expressed in SLC, and ACAN and LOXL2 in MLC. Regeneration factors also showed distinct expression patterns, with SOX9 and NT5E being highly expressed in MLC, and CXCL12, PDGFRB, ENG, and RUNX2 in SLC.

The susceptibility of MLC to IL-1β treatment was evident from the SEM analysis, which showed severe ECM degradation in IL-1β treated MLC compared to that in SLC. Transcriptomic profiling after post-IL-1β treatment revealed significant overexpression of OA development genes, such as MMP13, POSTN, and ADAMTS5, with higher levels in MLC. Cartilage regeneration genes such as SOX9, MCAM, and ACAN were downregulated in IL-1β treated samples, correlating with the observed phenotypic degradation.

To validate our *in vitro* goat MLC model, we compared IL-1β induced changes in the *in vitro* goat TMJ-OA model with rabbit TMJ-OA models(Tosa et al., 2023). Reanalysis of the bulk RNA-seq data from rabbit TMJ-OA SZ and CC, representing SLC and MLC, respectively, revealed consistent gene expression patterns. IL-1β treated goat SLC showed consistent up- and down-regulation of genes, while a large number of overexpressed genes in MLC correlated with the IL-1β treated goat MLC profile. GSEA of rabbit TMJ-OA MLC highlighted the enrichment of chondrocyte apoptosis-and inflammatory immune response-related gene sets, including mast cell and macrophage activation, cytokine production, neuroinflammatory responses, and elevated production of IL-1 and IL-6.

Further analysis identified OA-associated transcription factors (TFs), senescence genes, and epigenetic regulators in IL-1β treated goat MLC. Significant overexpression of OA-associated TFs, such as ADAMTS7(Liu et al., 2006; Zhang et al., 2015) and NOV(Kuwahara et al., 2020) has been observed, contributes to cartilage degradation and inflammation.

Cellular senescence factors including CSF2(Shin et al., 2023), ICAM1(Law et al., 2020) and CXCL16(van der Voort et al., 2005), were markedly overexpressed, confirming the specificity of the *in vitro* model for TMJ-OA studies. Additionally, epigenetic regulators such as ARID3A(Ratliff et al., 2019) and ARID5A(Masuda et al., 2013) are overexpressed, contributing to OA pathogenesis. A limitation of this study is the lack of human TMJ transcriptional signatures for comparison. Nevertheless, our findings provide valuable insights into the unique differentiation of the TMJ cartilage and its potential as a therapeutic target. TMJ disorders affect a significant proportion of the population and have limited treatment options, making the development of new therapies crucial for improving patient outcomes.

In conclusion, our study demonstrated the unique transcriptomic signature of the TMJ compared with the knee joint, highlighting its vulnerability to OA and pain. The distinct molecular profiles of SLC and MLC within the TMJ further emphasize the complexity of the cartilage regeneration and degeneration processes. The IL-1β induced OA-like changes in the MLC layers correlated with the rabbit TMJ-OA model, validating the *in vitro* goat TMJ-OA model. Hence, our study identified that IL-1β induced OA-like changes in goat TMJ MLC. This model could be useful for screening potential anabolic agents to evaluate the efficacy and mechanism before testing in large animal models of TMJ-OA. These findings provide valuable insights into the molecular mechanisms underlying TMJ disorders, and potential therapeutic targets for OA treatment.

## Support

This study was supported by an NIH grant, R01 DE031413 (MB).

## Ethical Compliance

All procedures performed in this study involving human participants were in accordance with the ethical standards of the institutional and/or national research committee.

## Author Contributions

The specific contributions are as follow. The conception and design of the study: Rajnikant Dilip Raut, Pushkar Mehra, Alejandro Almarza, Harpreet Singh, Louis Gerstenfeld, and Manish V. Bais. Acquisition of data, its analysis and interpretation: Rajnikant Dilip Raut, Chumki Choudhury, Amit Kumar Chakraborty, and Manish V. Bais. The drafting the article: Rajnikant Dilip Raut, Pushkar Mehra, Alejandro Almarza, Harpreet Singh, Louis Gerstenfeld, and Manish V. Bais. Revision of the manuscript for important intellectual content: Louis Gerstenfeld, Alejandro Almarza and Manish V. Bais. All the authors approve of the final version of the manuscript.

## Conflict of interest

The authors declare that they have no conflicts of interest regarding the content of this manuscript.

## Financial Interests

The authors declare that they have no financial interests regarding the content of this manuscript.

## Notes

### Competing Interest Statement

The authors have declared no competing interest.

